# GVRP: Genome Variant Refinement Pipeline for variant analysis in non-human species using machine learning

**DOI:** 10.1101/2024.01.14.575595

**Authors:** Jeonghoon Choi, Bo Zhou, Giltae Song

**Affiliations:** Division of Artificial Intelligence, School of Computer Science and Engineering, Pusan National University, Busan, South Korea; Department of Psychiatry and Behavioral Sciences, Stanford University School of Medicine, Stanford, CA, USA

**Keywords:** Genomic Variant, Non-Human Species, Machine Learning

## Abstract

Many investigations of human disease require model systems such as non-human primates and their associated genome analyses. While DeepVariant excels in calling human genetic variations, its reliance on calibrating against known variants from previous population studies poses challenges for non-human species.

To address this limitation, we introduce the Genome Variant Refinement Pipeline (GVRP), employing a machine learning-based approach to refine variant calls in non-human species. Rather than training separate variant callers for each species, we employ a machine learning model to accurately identify variations and filter out false positives from DeepVariant.

In GVRP, we omit certain DeepVariant preprocessing steps and leverage the ground-truth Genome In A Bottle (GIAB) variant calls to train the machine learning model for non-human species genome variant refinement. We anticipate that GVRP will significantly expedite genome variation studies for non-human species,.

## Introduction

The swift advancements in genome sequencing technologies have spurred many explorations into the complexities of genome variations and their functional links to diseases, including cancers and developmental disorders. The experimental investigations of these associations often involve the utilization of non-human model organisms such as non-human primates [1,2] and mice [3-5].

The conventional approach to genomic variant identification entails aligning sequencing reads with a well-established reference genome, e.g. GRCh38, and then determining the discrepancies between the reads and the reference sequence. To achieve precise variant detection, indispensable preprocessing steps are employed to eliminate noise using known information. While the widely adopted Genome Analysis Toolkit (GATK) [6] excels in offering comprehensive preprocessing and variant calling functionalities, its exclusive reliance on genome sequence information limits its variant calling accuracy when compared to more sophisticated methods rooted in machine learning and deep learning.

Machine learning and deep learning, key components of artificial intelligence, have revolutionized various domains, driven by advancements in computational power and the availability of extensive datasets. These technologies go beyond enhancing existing methods; they are unlocking new insights into genomic complexity. Their advancements have thus opened new possibilities for accurate and efficient genome variant analysis, positioning them at the forefront of genomic innovation.

Various genome variant refinement filters based on machine learning and deep learning have been explored to improve upon traditional statistical-based variant callers, including SNooPer [7], RFcaller [8], and VEF [9], which employ ensemble decision tree techniques [10-12], and utilizing logistic regression [13] to filtering false positive variant call [14]. SNooPer employs Fisher’s exact test to learn the quality scores of called variants, thereby filtering out false positives. Similarly, RFcaller uses read information, quality scores, and coverage data from variants called by bcftools [15] to filter false positive variant calls. Both VEF and logistic regression-based methods refine the process by utilizing quality scores and read information from variants called by GATK. While these refinement approaches enhance the performance of conventional statistical variant callers and improve the quality of variant calling, they still have limitations due to the inherent accuracy constraints of the existing variant callers.

Deep learning has been employed to make fundamental improvements in variant call quality. A notable example of which is DeepVariant [16]. DeepVariant encodes read bases, quality scores, and other read features from alignments into RGB pile-up images, which are then used to train a convolutional neural network (CNN) [17]. This process enables the calling of single nucleotide polymorphism (SNP) and insertion & deletion (INDEL). Similar models include DeepTrio [18] and DeNovoCNN [19], which also utilize pile-up images and train them with CNNs. However, these models incorporate additional features for encoding or using Deep Neural Networks [20] to call *de novo* SNPs and INDELs specifically. These deep learning-based variant callers have demonstrated superior performance compared to traditional statistical-based callers, with DeepVariant standing out for its exceptional performance among various deep learning-based callers [21-23].

The applicability of methods such as DeepVariant to non-human species is constrained by certain limitations. In the preprocessing of sequencing data, DeepVariant relies on calibration against previously known genomic variants, most commonly compiled from human population studies such as the 1000 Genomes Project. For non-human species, such types of genome variants are not available, thus impeding the seamless application of these preprocessing steps. Furthermore, DeepVariant, initially trained on human genotypes, exhibits diminished performance when applied to non-human species (given its model training is human-specific). Recognizing this limitation, attempts have been made to broaden DeepVariant’s utility across non-human species. An illustrative example is a study [24] wherein DeepVariant underwent re-training for the mosquito genome. However, it is worth noting that retraining DeepVariant for each non-human species genome poses a considerable challenge and not practical.

To address these challenges, we introduce the Genome Variant Refinement Pipeline (GVRP), a machine learning-based approach for refining variant calls in non-human species genome analysis. Rather than training a separate variant caller for each species, GVRP is an end-to-end pipeline capable of refining variant calls for genomes of any non-human species.

## Methods

### Non-human Species Genome Sequence Data Preprocessing

Since population-derived genomic variants are usually not available for non-human species, the variant calling procedure using DeepVariant is different for human and non-human species (Figure 1). To assess the effectiveness of DeepVariant on non-human species as a baseline for GVRP development, we first process two human genomes with known ground-truth variants using the same approach as one would for non-human species and then compare the variants determined in this manner against their known ground-truths. These two human genomes are NA12878/HG001 [25] and HG002 [26] (part of the Ashkenazi Trio) where their respective high-quality ground-truth variants have been extensively and thoroughly determined as a standard for genome analysis through the Genome in a Bottle (GIAB) consortium.

**Figure 1.**
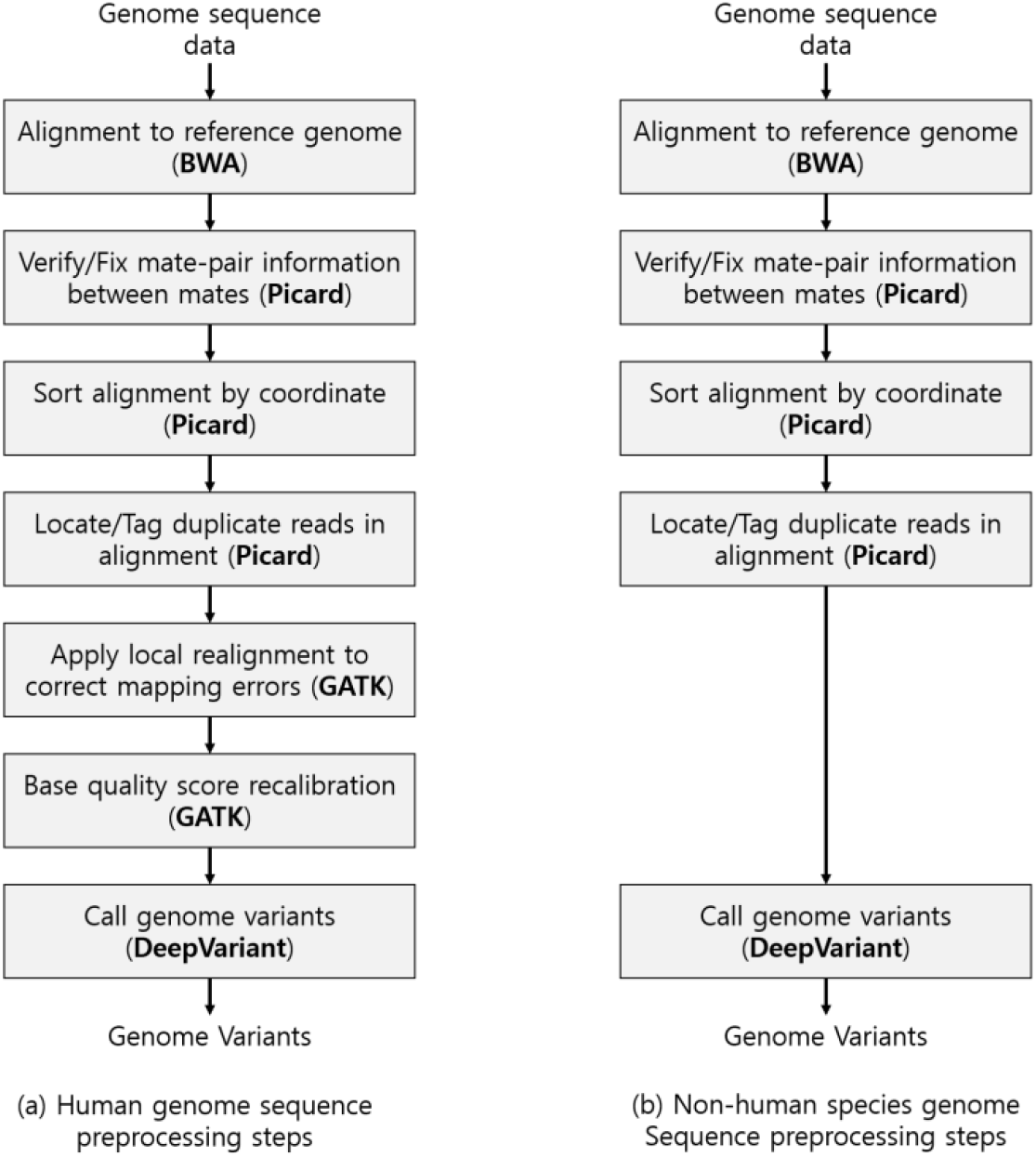
Comparison of the variant calling workflows for human and non-human genomes. For human genomes (a), the sequence reads are aligned to a reference genome and undergo preprocessing steps like INDEL realignment and base quality score recalibration with GATK, before using DeepVariant to call variants. For non-human genomes (b), the workflow must bypass these preprocessing steps, proceeding directly from read alignment and duplicate tagging with Picard to variant calling with DeepVariant, as steps such as local realignment require known variants determined *a priori* from population studies.

To call variants with DeepVariant as if the human genome sequencing data for HG001/NA12878 and HG002 were those of non-human species, we must bypass the steps of INDEL realignment and base recalibration (Fig. 1 (b)). Both steps require knowledge of pre-existing variants, which is usually not available for non-human species due to less extensive population-scale studies. Normally, these steps are critical for variant calling in human genomes to ensure data quality [24] (Fig. 1). INDEL realignment corrects the alignment of sequences around insertions and deletions, where mapping is typically difficult and requires a database of known indels for guidance. Base recalibration adjusts base quality scores to account for systematic errors of sequencing machines and relies on known variant information for accuracy [24].

### Non-Human Genome Sequence Preprocessing for Human Genome Sequence

We used BWA-MEM [27] v. 0.7.17 to align HG001 and HG002 paired-end sequencing data (∼60x coverage) to the GRCh38 [28]. To fix mate information, sort alignment, and mark duplicates, we used FixMateInformation, SortSam, and MarkDuplicates from Picard [29] v. 3.1.0 respectively. We used ‘True’ for both the ‘ADD_MATE_CIGAR” and ‘ASSUME_SORTED’ options and ‘SILENT’ for ‘VALIDATION_STRINGENCY’ option for all 3 processes. We used samtools [30] v.1.3.1 for indexing the alignment and DeepVariant v. 1.4.0 to obtain variant call (SNPs and INDELs) in Variant Call Format (VCF) format.

### DeepVariant

Google’s DeepVariant is a leading variant caller that outperforms other existing tools in identifying variants in the human species [31]. It uses Convolution Neural Network (CNN) [32] and known genotypes for analyzing genome sequencing data. First, DeepVariant encodes the reads bases, quality scores and other read features into a red-green-blue pileup image at a particular variant site. The CNN model is then trained using the pileup images of known genotypes. Once trained, the CNN can predict genotype probabilities for each site. Finally, DeepVariant determines variant calls based on the most likely genotype, whether heterozygous or homozygous non-reference. Since DeepVariant is trained on human genotypes, validation of its performance for non-human genome sequence data is critical.

### Refinement Pipeline for The Non-Human Species Variant Call from DeepVariant

To refine DeepVariant call for non-human species, we developed Genome Variant Refinement Pipeline (GVRP). GVRP refines variant call ***V*** from DeepVariant to refined variant call ***V***^*^ by pre-trained machine learning model ***M*** (Fig. 2). DeepVariant’s variant call ***V*** is a VCF file that that contains meta-information and the information for each specific variant call, *υ*, which includes details such as the chromosome (**CHROM**), position (**POS**), reference base (**REF**), alternate base (**ALT**), phred-scaled quality score (**QUAL**), and the genotype field. Genotype field presents the genotype information about *υ*, there are 6 features: genotype (**GT**), unfiltered allele depth (**AD**), read depth at this position (**DP**), conditional genotype quality (**GQ**), phred-scaled phenotype likelihood (**PL**) and variant allele fraction (**VAF**).

**Fig. 2.**
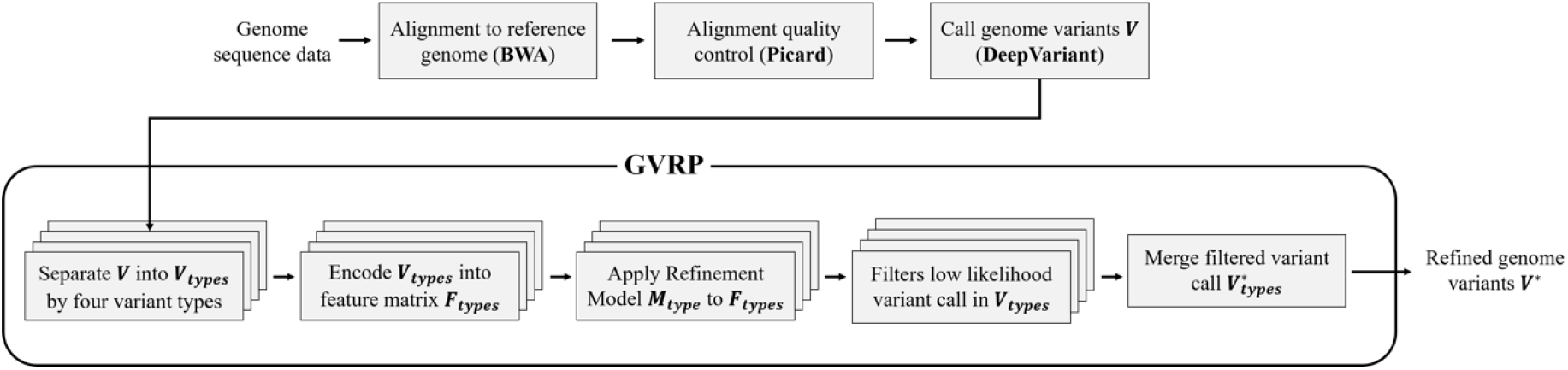
Non-human species variant call process with GVRP. Initially, the GVRP takes as input the variant call ***V*** from DeepVariant and separates them into distinct variant types, denoted as ***V***_*types*_. Subsequently, ***V***_*types*_ are encoded into a feature matrix *F*_*types*_ utilizing genotype information and quality scores. Upon encoding to *F*_*types*_, GVRP applies the refinement model ***M***_*types*_ for each respective variant type, which predict the accuracy likelihood of the variant calls and filter out those with low likelihood within ***V***_*types*_. Finally, GVRP concatenates the refined variant calls 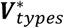 into the refined set ***V***^*^.

### Separate Variant Call in Four Types

GVRP first separates ***V*** into four ***V***_*types*_ by the type of variant. ***V*** can be categorized in two categories, whether the variant occurs in heterozygous or homozygous, and variant effects only in SNP or not. In **GT**, allele values are separated by / or |, 0 for the reference allele, 1 and 2 for each allele listed in **ALT**. For the **REF** and **ALT**, the reference base(s) and alternate base(s) are recorded. We can easily separate ***V*** into four types: Heterozygous-INDEL (***V***_*HT−INDEL*_), Heterozygous-SNP (***V***_*HT−SNP*_), Homozygous-INDEL (***V***_*HM−INDEL*_), and Homozygous-SNP (***V***_*HM−SNP*_), by using this information. A detailed description of each type per condition of **GT** and **REF**/**ALT** is shown in Table 2.

**Table 1.** Genome sequence preprocessing parameter.

| Toolkit | Command | Usage | Parameter | value |
| --- | --- | --- | --- | --- |
| BWA | MEM | Alignment | -k | 100000000 |
| Picard | SortSam | Sort alignment by coordinate | ADD_MATE_CIGAR<br>ASSUEM_SORTED<br>VALIDATION_STRINGENCY | TRUE |
|  | FixMateInformation | Verify mate-pair information |  |  |
|  | MarkDuplicates | Locates and tags duplicate reads |  |  |

**Table 2.** Conditions of **GT, REF**, and **ALT** for four variant types: Heterozygous-INDEL, Heterozygous-SNP, Homozygous-INDEL, and Homozygous-SNP.

|  | Separated values on **GT** | Length of base(s) in **REF** and **ALT** | Recorded base(s) in **REF** and **ALT** |
| --- | --- | --- | --- |
| $V_{HT-INDEL}$ | Different | Different | Subset of another |
| $V_{HT-SNP}$ | Different | Same as 1 | Different |
| $V_{HM-INDEL}$ | Same | Different | Subset of another |
| $V_{HM-SNP}$ | Same | Same as 1 | Different |

### Learning The Refinement Models for Each Type

After separating ***V***_*types*_ by the type of variant, GVRP makes feature matrix *F*_*types*_ from ***V***_*types*_ for each type to predict the validity of each variant. We used **QUAL, AD, DP, GQ, PL**, and **VAF** information from ***V***_*types*_ to make the feature matrix for learning/predicting variant call. Unlike **QUAL, DP**, and **GQ**, which record scalar values, for **AD, PL**, and **VAF**, the record is formed as a list of values. We merged the list of values by mean and the difference between max and min values. To labeling call for learning, we matched the *υ* to GIAB-published ground-truth variants using **CHROM** and **POS** as the key. For the variant, which is not in the ground-truth or different **REF, ALT**, and **GT** information are labeled as “miscalled”. This labeling step only processes for learning refinement model ***M***.

Finally, GVRP applies ***M***_*types*_ to *F*_*types*_ per type, respectively, and predicts the quality of the variant. We trained various machine learning models widely used in diverse fields, including the tree-based models such as LGBM [33], XGBoost (XGB) [34], Random Forest (RF), as well as traditional machine learning methods such as logistic regression (LR), k-nearest neighbor (k-NN) [35] classifier, and naïve Bayes classifier (NB) [36] for each type. GVRP predicts false positives and delete them from each ***V***_*types*_ and the results for each 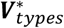 are then merged into ***V***^*^, ensuring that the elimination of the False-Positive variants in original ***V***.

## Results

### The performance of DeepVariant on preprocessed human genome sequencing data, similar to non-human species

Table 3 presents the performance of DeepVariant when identifying variants from HG001 and HG002 sequencing data that were processed like non-human species when compared against the GIAB ground-truth variants. For HG002, DeepVariant identified a total of 4,694,623 variants, out of which 777,192 were miscalled, resulting in a miscall rate of 16.56%. In the case of HG001, the miscall rate was 19.58%. These figures suggest that when DeepVariant operates without two preprocessing steps that are specific to human genomic data, its accuracy significantly drops. Therefore, for non-human genomes, DeepVariant requires a specialized refinement model to improve variant calling accuracy.

**Table 3.**
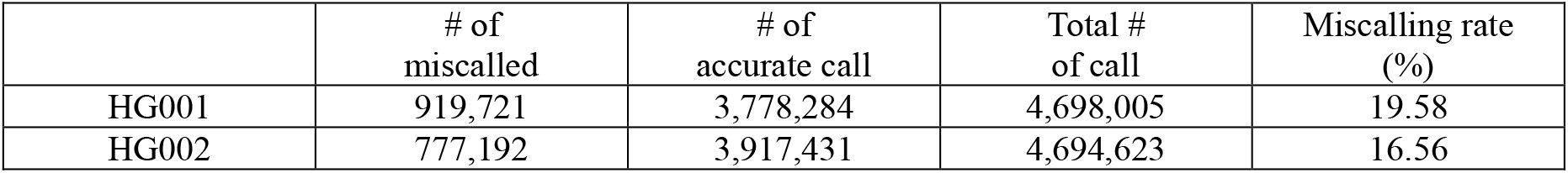
DeepVariant performance on human sequencing data preprocessed similarly to non-human species.

|  | # of<br>miscalled | # of<br>accurate call | Total #<br>of call | Miscalling rate<br>(%) |
| --- | --- | --- | --- | --- |
| HG001 | 919,721 | 3,778,284 | 4,698,005 | 19.58 |
| HG002 | 777,192 | 3,917,431 | 4,694,623 | 16.56 |

### Validation Dataset – Human Genome Sequence

To evaluate GVRP, we used high-quality genome sequence data that had previously been used in DeepVariant. Specifically, we made use of the HG001 and HG002 genome datasets, which had been used in the evaluation of DeepVariant. This approach is suitable for evaluating our model as these two datasets are independent of each other. Furthermore, the ancestries of these individuals are different; HG001 is of European (from the CEPH Utah Reference Collection), while HG002 is of Ashkenazi Jewish ancestry.

We evaluate our pipeline for learning on the HG001 dataset and validating on the HG002 dataset, and vice versa. We trained a machine learning model for each variant type. The number of variant calls from DeepVaraint for HG001 and HG002 are shown in Table 4 and Table 5, respectively. We compare various machine learning models and simple deep neural network model for the refinement model.

**Table 4.**
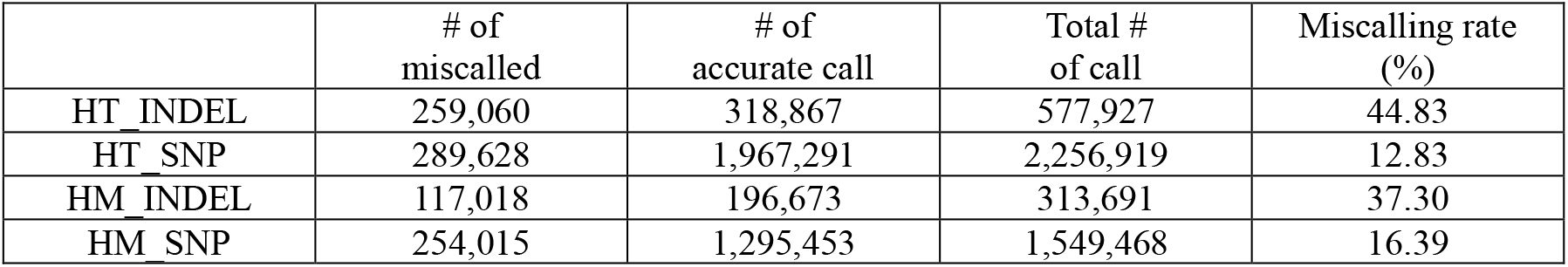
The number of variants for each type in HG001.

**Table 5.** The number of variants for each type in HG002.

|  | # of<br>miscalled | # of<br>accurate call | Total #<br>of call | Miscalling rate<br>(%) |
| --- | --- | --- | --- | --- |
| HT INDEL | 216,936 | 354,699 | 570,635 | 37.84 |
| HT SNP | 184,510 | 2,048,859 | 2,233,369 | 8.26 |
| HM INDEL | 113,999 | 208,758 | 322,757 | 35.32 |
| HM SNP | 262,747 | 1,305,115 | 1,567,862 | 16.76 |

### Validation for Human Genome Sequence Datasets

We validated various machine learning models and simple deep neural network for non-human species like preprocessed human genome sequence data. Displayed in Tables 5, each model’s performance is quantified across four distinct data types, using the HG001 and HG002 datasets for both training and evaluation steps. Our refinement model excels at correctly classifying accurate and miscalled variant calls. Tables 5 and 6 show the merged performance of model on each variant type. We can observe that the machine learning-based variant refinement models perform well in classifying accurate variants and identifying miscalled variants. Additionally, the tree-based boosting model and k-NN model exhibit superior performance compared to other models, achieving F1 scores of 0.932 and 0.948, respectively, for XGBoost (XGB), LGBM, and k-NN trained on HG002/HG001.

**Table 6.**
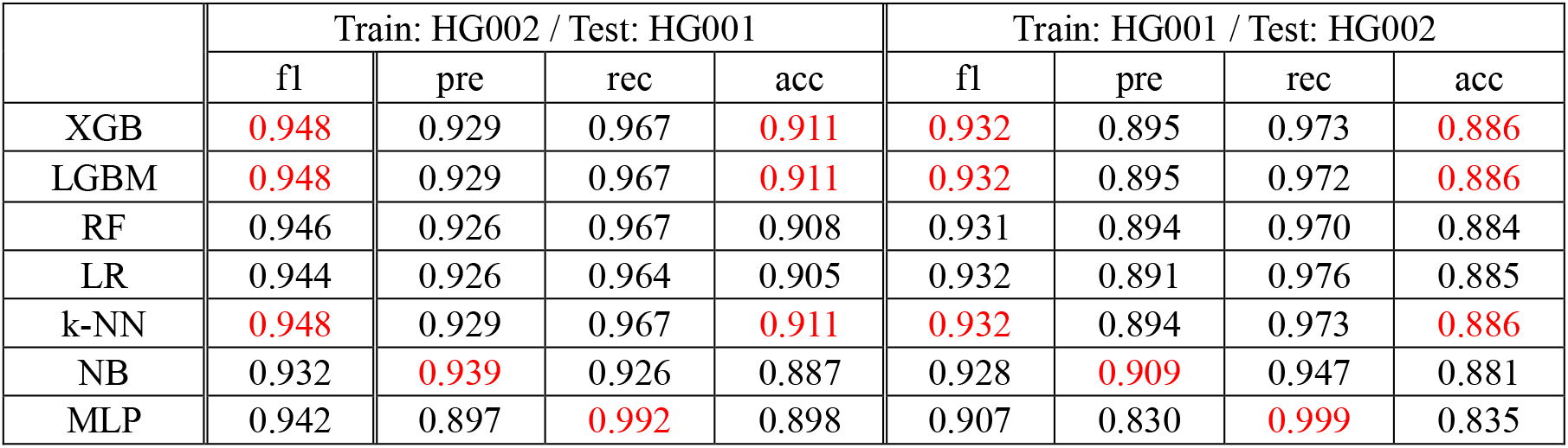
Evaluate HG001/HG002 trained refinement model on HG002/HG001.

For model evaluation on each variant type, the model for SNP variant type performs more better than INDEL variant type (Fig. 3). For HG002 genome variant trained refinement model evaluate on HG001, the highest F1 score achieved for HT-INDEL, HT-SNP, HM-INDEL, and HM-SNP were 0.838, 0.951, 0.852, and 0.944, respectively. In a reverse scenario, where training occurred on HG001 and validated on HG002, the best F1 scores were 0.867, 0.974, 0.865, and 0.945 for the respective categories. As the results show, the INDEL type refinement model performs at least 9.24% better than the SNP refinement model. The performance difference between INDEL and SNP arises from the significantly larger volume of INDEL variant data, despite the imbalance in the INDEL variant data. Furthermore, our refinement models exhibited stable performance (not exceeding 3%) across these two genomic datasets.

**Figure 3.**
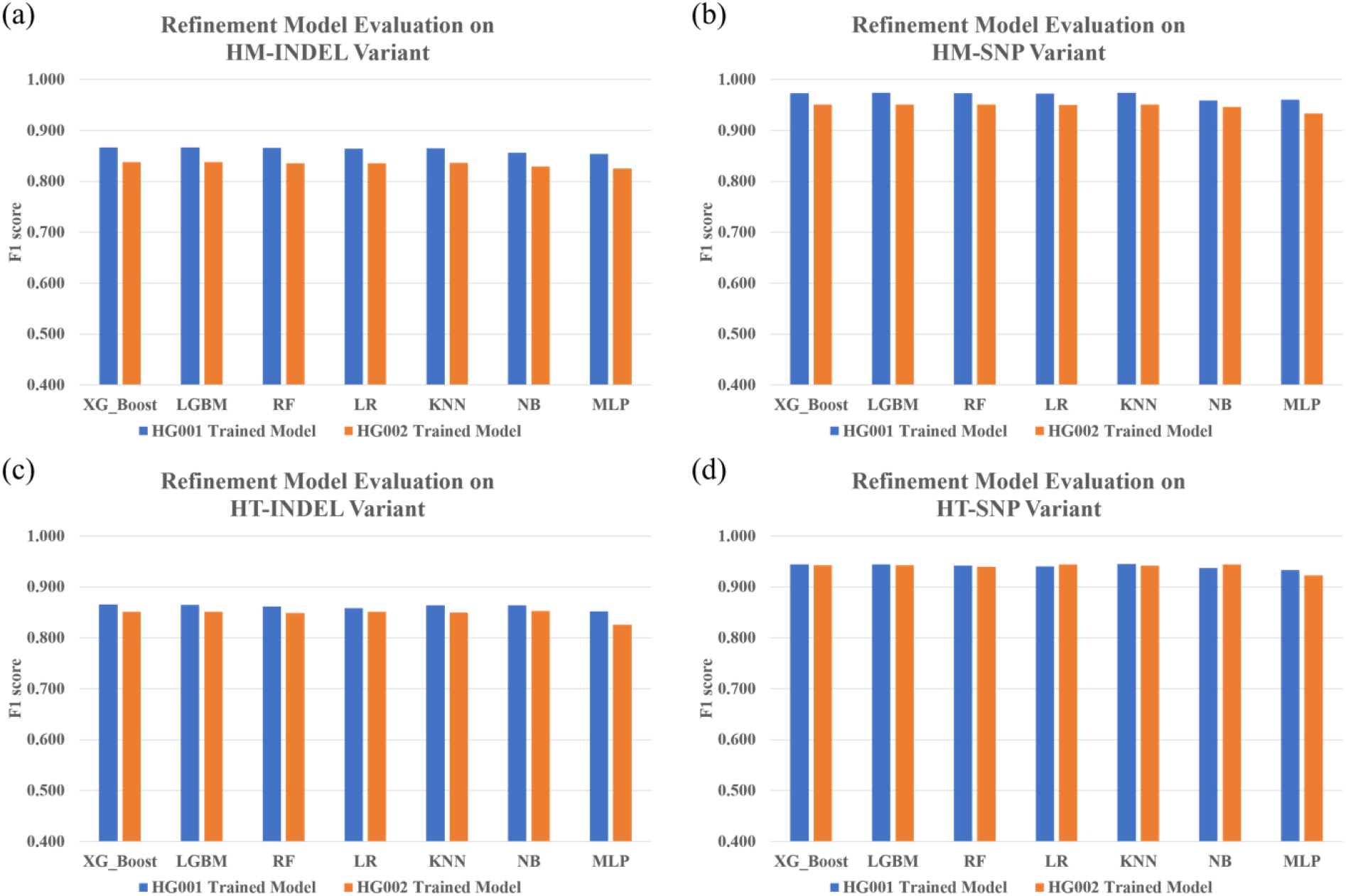
presents a comparative analysis of F1 scores across four types of refinement models, each trained on distinct variant datasets: HM-INDEL (a), HM-SNP (b), HT-INDEL (c), and HT-SNP (d). The blue bars represent the refinement models trained on the HG001 dataset, while the orange bars indicate those trained on the HG002 dataset. It is observed that the F1 scores for SNP refinement models are consistently above 0.9, in contrast to the INDEL models, which score below 0.9. The performance of the models is notably consistent between the HG001 and HG002 genomic variant datasets, suggesting stable model behavior across different genomic data.

## Conclusion

In this study, we developed the Genome Variant Refinement Pipeline (GVRP), a machine learning-based pipeline designed to enhance the accuracy of variant calling for non-human species by DeepVariant. This paper highlights two main findings: firstly, DeepVariant’s limitations when applied to non-human species genome data, and secondly, the application of machine learning to refine variant calls in non-human genome data. For validation and model training purposes, we utilized HG001 and HG002 datasets from the GIAB consortium. By omitting INDEL realignment and base recalibration, we processed them as one would for non-human species data (where population-derived known variants are usually not available). This revealed that DeepVariant produced a higher rate of miscalls for genomes processed in this manner than for those with standard DeepVariant preprocessing procedures. To overcome this, we developed a pipeline that filters out false positives. For more precise model training we categorized variant calls into four groups based on size and position (genotype?). Our refinement model demonstrated its effectiveness in correcting miscalled variants from the sequencing of non-human species. We note that GVRP has not yet been directly tested on actual non-human species. While our machine learning model outperformed basic DNN approaches, there is potential for further enhancement using cutting-edge deep learning models. In our future work, we plan to test our model on real non-human species data and enhance the refinement model with state-of-the-art deep learning techniques.

